# Anxiety and risk-taking behavior maps onto opioid and alcohol polysubstance consumption patterns in male and female mice

**DOI:** 10.1101/2024.08.22.609245

**Authors:** Makenzie Patarino, Ziheng Christina Wang, Andrew Byungwook Kim, Katrina Wong, Suhjung Janet Lee, Emma Skillen, Richa Nag, Britahny Baskin, Abigail G. Schindler

## Abstract

Polysubstance use is prevalent in the population but remains understudied in preclinical models. Alcohol and opioid polysubstance use is associated with negative outcomes, worse treatment prognosis, and higher overdose risk; but underlying mechanisms are still being uncovered. Examining factors that motivate use of one substance over another in different contexts in preclinical models will better our understanding of polysubstance use and improve translational value. Here we assessed baseline anxiety-like and locomotive behavior and then measured voluntary consumption of multiple doses of alcohol and fentanyl in group housed male and female mice using our novel Socially Integrated Polysubstance (SIP) system. Fifty-six male (n=32) and female (n=24) adult mice were housed in groups of 4 for one week with continuous access to food, water, two doses of ethanol (5% and 10%) and two doses of fentanyl (5 ug/ml and 20 ug/ml). Our analyses revealed sex differences across multiple domains – female mice consumed more liquid in the dark cycle, had higher activity, a higher preference for both ethanol and fentanyl over water, and their fentanyl preference increased over the seven days. Furthermore, both male and female mice displayed polysubstance consumption patterns, with female mice displaying more prolonged polysubstance use across days in the SIP chambers. We then used machine-learning techniques to reveal underlying relationships between baseline behavioral phenotypes and subsequent polysubstance consumption patterns, where anxiety- and risk-taking-like behavioral phenotypes mapped onto discrete patterns of polysubstance use, preference, and escalation. By simulating more translationally relevant substance use and improving our understanding of the motivations for different patterns of consumption, this study contributes to the developing preclinical literature on polysubstance use with the goal of facilitating better treatment outcomes and novel therapeutic strategies.

## Introduction

Polysubstance use, or the longitudinal, sequential, or simultaneous use of multiple substances, is a persistent and growing concern globally. Clinical populations that engage in polysubstance use experience detrimental outcomes including worsened substance use disorder (SUD) severity, mental and physical health status, treatment response, and mortality, as well as increased risk for overdose, suicide, and infectious and sexually transmitted disease^1,2^. In one longitudinal study, persistent polysubstance use was associated with the poorest biological aging and midlife health and financial/social preparedness^3^. Nearly all individuals with a SUD additionally consume other substances and the majority have at least one other diagnosed SUD^4^.

The increasing prevalence of alcohol and opioid co-use is a pressing concern – from 2002 to 2012, there was a 15-fold increase in the number of individuals with AUD and comorbid OUD^5^. Alcohol and opioid co-use accelerates the progression of problematic use and is more harmful than either substance used alone^6^ (the number of deaths resulting from opioid overdose also involving alcohol increased 5.5 times between 1999 and 2017^7^). Thus, there is an urgent need to address alcohol and opioid polysubstance use to limit harms and improve outcomes.

The choice to use one or more substances may depend on life history, current environment, and personality type. Experiencing stressful life events is predictive of polysubstance use^8–13^. Additionally, maladaptive coping (including aggressive, reactive, or substance-driven coping) is thought to play a role in mediating polysubstance use^14–16^. Behavioral or psychological phenotypes of an individual may also influence which substance or substances to use, and if that choice remains constant in all cases or is circumstance dependent. For example, both preclinical and clinical studies have shown that higher levels of anxiety and novelty-seeking are correlated with increased alcohol consumption, albeit with different patterns of use^17–19^. There may be a relationship between anxiety-like or reward behaviors and increased opioid consumption, but findings remain mixed^20,21^. However, these types of investigations have been limited to a single substance, so conclusions about pre-existing behaviors or personality traits and polysubstance use remain limited. Sex differences in substance use are also thought to be a critical factor, but little focus has been placed on understanding polysubstance use in relation to biological sex.

While work to further characterize polysubstance use patterns in clinical populations is ongoing, preclinical models present a viable line of research to investigate underlying motivations and mechanisms. Preclinical polysubstance use research typically involves alcohol, nicotine, or cocaine, with limited studies on cannabinoids, hallucinogens, and opioids. Within preclinical opioid research, heroin is commonly administered over prescription opioids or fentanyl. Even though alcohol and opioid co-use is quite common, animal studies involving the combination of these two substances are lacking. Furthermore, many current studies lack additional features of realistic human substance use, such as a group-housed social environment during use and voluntary, continuous access to multiple substances and concentrations.

To address these gaps, the current study investigated voluntary intake of alcohol, fentanyl, and water in a group-housed environment in adult male and female mice. To do this, we utilized the Socially Integrated Polysubstance (SIP) system, which allows rodents to remain group-housed while self-administering substances with continuous monitoring and intake measurement^22^. Previous research using SIP cages in our lab revealed differences in activity and flavor preference between male and female rodents, offering insights into how sex may influence substance preference and behavior patterns.

## Materials and Methods

### Animals

All experiments utilized female and male (as determined by genital appearance at weaning) C57BL/6 mice from Jackson Labs aged 9-11 weeks of age at time of arrival to VA Puget Sound. Mice were housed by sex in cages of four on a 12:12 light:dark cycle (lights on at 06:00), and were given ad libitum food and water. All animal experiments were carried out in accordance with AAALAC guidelines and were approved by the VA Puget Sound IACUC. Mice were acclimated to the VA for one week following arrival and subsequently handled for an additional week prior to experimental manipulation. To increase rigor and reproducibility, the study included at least two cohorts of mice each run at separate times.

### Baseline behavioral testing

One week prior to housing in the SIP cages, animals were tested in the open field and then at least 24 hours later in the elevated zero maze to assess locomotion and anxiety-like behavior. On each day of testing, animals were allowed at least 30 minutes to acclimate to the testing room.

#### Open field box (OFB)

Mice were allowed 5 minutes to explore a large circular open space (1 meter diameter) and their movements were recorded from above and analyzed with Anymaze (Wood Dale, IL). Decreased time spent in the middle of the OFB is indicative of an anxiety-like phenotype.

#### Elevated zero maze (EZM)

Mice were allowed 5 minutes to explore an elevated zero maze (Maze Engineers, Skokie, IL) and their movements were recorded from above and analyzed with Anymaze (Wood Dale, IL). Decreased time spent exploring the open arms is thought to reflect anxiety-like behavior.

### RFID transponder implantation

Each RFID transponder (Euro I.D., Koln, Germany) is coated in a biocompatible glass material and is 2.12 mm x 12 mm diameter. At least 72 hours prior to SIP cage housing, each transponder is sterilized and injected subcutaneously behind the shoulder blades of an anesthetized mouse (5% isoflurane) using the provided syringe applicator.

### Socially Integrated Polysubstance (SIP) system

As previously described, the SIP system enables group housed mice to self-administer multiple different substances in a home-cage setting while still maintaining individual intake levels on a second-to-second time scale (Wong et al., 2023). The current study employed a setup with six drinking stations in a rectangular home cage design (3 drinking stations on each long wall). Mice were housed for seven days with continuous access to water (2 drinking stations, one on each wall), two different doses of ethanol (5% and 10%) and two different doses of fentanyl (5 ug/ml and 20 ug/ml). Cages were checked daily, and food was available ad libidum. Custom Python scripts were used to integrate the RFID and VDM data streams via common timestamps.

### Unsupervised machine learning (cluster analysis) of baseline behavioral testing

Given the high degree of collinearity across the 12 behavioral parameters collected from the OFB and EZM, we first performed a dimensionality reduction step using Principal Component Analysis (PCA). We then used the first three principal components (explaining over 75% of model variance) in a K-means cluster-based approach. Cluster stability was assessed as previously described^23^, using the scores for homogeneity, adjusted Rand, and adjusted mutual information criterion and a bootstrap approach with repeated random assignment of initial cluster centroids. K=3 clusters was chosen based on the above evaluation metrics.

### Statistical analysis

Data are expressed as mean ± SEM. Differences between groups were determined using a two-tailed Student’s t-test, one-way analysis of variance (ANOVA), or two-way (repeated measures when appropriate) ANOVA followed by post hoc testing using Bonferroni’s Multiple Comparison. Reported p values denote two-tailed probabilities of p ≤ 0.05 and non-significance (n.s.) indicates p > 0.05. Statistical analysis and visualization were conducted using Graph Pad Prism 9.0 (GraphPad Software, Inc., La Jolla, CA) and with custom Python scripts.

## Results

Fifty-six male (n=32) and female (n=24) adult mice were housed in groups of 4 for one week in the Socially Integrated Polysubstance (SIP) system with continuous access to water, two doses of ethanol (5% and 10%) and two doses of fentanyl (5 ug/ml and 20 ug/ml). Visits to the drinking chambers were collected at 100Hz and drinking data was collected at 1Hz. Across the 56 mice, the data set included over 650,000 RFID data points and over 45,000 drinking data points.

Data was first examined by grouping all substances together to assess general differences in the amount of time spent in the drinking chambers and total intake male and female mice (Supp Figure 1). Female mice spent more time in the drinking chambers than male mice (Supp Figure 1a-e) but did not differ in total amount of fluid consumed (Supp Figure 1f-j).

Next we examined potential sex differences in fluid intake for each substance separately (Figure 1). When summarized across all days, there was a significant interaction effect (two-way RM ANOVA: interaction effect F[4,216]=4.6, p<0.001, main effect Sex F[1,54]=1.2, p>0.05, main effect Substance F[4,216]=23.7, p<0.0001; BMCT post hoc) (Figure 1a). Potential differences in amount consumed for males vs. females across substances and light/dark cycle was examined using a separate two-way ANOVA for light vs. dark cycle (Dark cycle: two-way RM ANOVA: interaction effect F[4,216]=4.6, p<0.001, main effect Sex F[1,54]=2.84, p>0.05, main effect Substance F[4,216]=24.6, p<0.0001; BMCT post hoc; Light cycle: two-way RM ANOVA: interaction effect F[4,216]=3.62, p<0.01, main effect Sex F[1,54]=0.38, p>0.05, main effect Substance F[4,216]=16.67, p<0.01; BMCT post hoc) (Figure 1b). Finally, we examined potential differences in male vs. female mice across days for each substance separately (Figure 1c; see Table 1 for statistical results). Heat maps depicting the amount of each type of substance consumed across individual mice are shown in Figure 1d. Heat maps depicting the average amount of each substance consumed for males and females across days and zeitgeber time are shown in Figure 1e.

**Figure 1:**
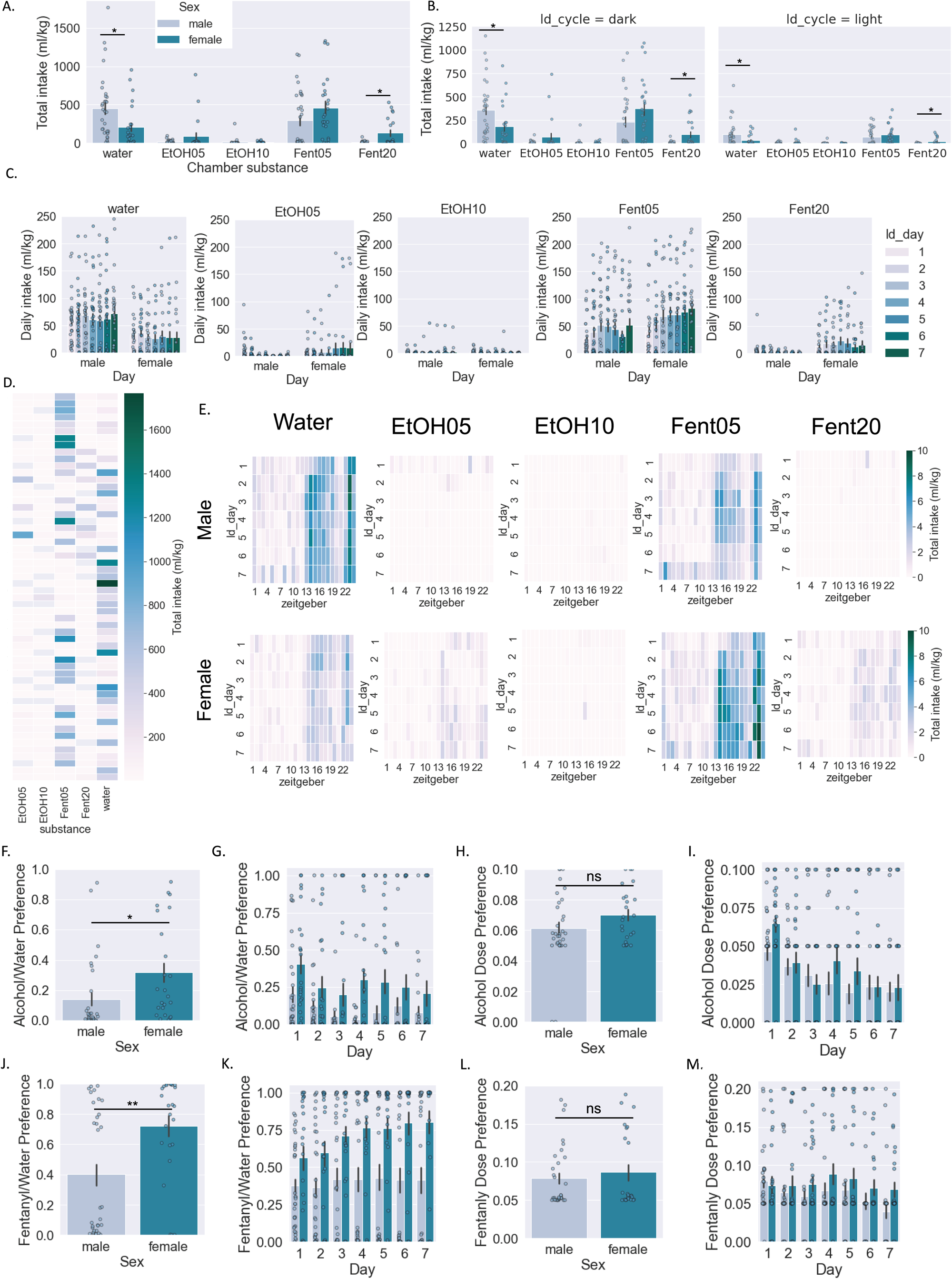
Consumption of individual substance/dose by sex. A-C: Amount of liquid consumed by male and female mice (normalized to body weight) for each individual substance/dose combination in total (a), across light/dark cycle (b), and across days (c). D. Heatmap of amount consumed for each substance for individual mice. E. Heatmap of amount consumed for each substance in male and female mice across days and Zeitgeber time. F-I: Alcohol preference over water (f,g) and alcohol dose preference (h,i). J-M: Fentanyl preference over water (j,k) and fentanyl dose preference (l,m). Two-way RM ANOVA *post hoc* BMCT (a-c, g,i,k,m). Student’s t-test (f,h,j,l). *p ≤ 0.05, **p ≤ 0.01. Values represent mean ± SEM.

**Table 1:** amount consumed across substance and day.

| Chamber | Interaction F(df) | Interaction P | Sex F(df) | Sex P | Day F(df) | Day P |
| --- | --- | --- | --- | --- | --- | --- |
| water | F[6,324] = 1.1 | 0.34 | F[1,54] = 5.9 | **0.02** | F[6,324] = 1.5 | 0.16 |
| etoh 5% | F[6,324] = 3.1 | **0.006** | F[1,54] = 2.8 | 0.1 | F[6,324] = 1.6 | 0.1 |
| etoh 10% | F[6,324] = 2.0 | 0.07 | F[1,54] = 0.11 | 0.74 | F[6,324] = 1.2 | 0.35 |
| fent 5ug | F[6,324] = 3.4 | **0.003** | F[1,54] = 2.5 | 0.12 | F[6,324] = 12.7 | **<.0001** |
| fent 20ug | F[6,324] = 2.0 | 0.07 | F[1,54] = 11.1 | **0.002** | F[6,324] = 1.0 | 0.45 |

When examining total intake, preference for both alcohol (Student’s unpaired t-test, t[54]=2.37, p<0.05) (Figure 1f) and fentanyl (Student’s unpaired t-test, t[54]=3.1, p<0.01) (Figure 1j) was higher in females. When examined across days, there were significant main effects of Sex and Day but no significant interaction effect for ethanol preference (two-way RM ANOVA: interaction effect F[6,324]=1.6, p>0.05, main effect Sex F[1,54]=8.2, p<0.01, main effect Day F[6,324]=1.8, p>0.05; BMCT post hoc) (Figure 1g). Conversely, when examining fentanyl preference across days, there was a significant interaction effect and significant main effects of Sex and Day (two-way RM ANOVA: interaction effect F[6,324]=2.9, p<0.01, main effect Sex F[1,54]=9.3, p<0.01, main effect Day F[6,324]=6.0, p<0.001; BMCT post hoc) (Figure 1k). Finally, we examined dose preference in males vs. females for alcohol (doses available were 5% and 10%) and fentanyl (doses available were 5 ug and 10 ug/ml) (Figure 1h-i,l-m). When examining total intake, dose preference for alcohol was not significantly different between males and females (Student’s unpaired t-test, t[54]=1.54, p>0.05) (Figure 1h) nor was dose preference for fentanyl (Student’s unpaired t-test, t[54]=0.67, p>0.05) (Figure 1l). When examined across days, there was only a significant main effect of Day but not Sex and no significant interaction effect for ethanol dose preference (two-way RM ANOVA: interaction effect F[6,168]=1.0, p>0.05, main effect Sex F[1,54]=1.7, p>0.05, main effect Day F[6,324]=6.5, p<0.001; BMCT post hoc) (Figure 1i). Likewise, when examining fentanyl dose preference across days, there was only a significant main effect of Day but not Sex and no significant interaction effect for ethanol dose preference (two-way RM ANOVA: interaction effect F[6,324]=1.0, p>0.05, main effect Sex F[1,54]=1.7, p>0.05, main effect Day F[6,324]=2.7, p<0.05; BMCT post hoc) (Figure 1m).

In addition to examining each substance separately, we also investigated potential polysubstance use in male and female mice (Figure 2). Here we classified polysubstance use as intake of multiple substances within a 60 minute time frame and visualized the data as nested pie charts. In each nested pie chart, the outer ring is the count of substance types consumed and the inner ring is the type of substance combination. We tested for potential significant differences using ci squared test and examined potential differences in the distribution of a) how many types of substances were consumed (substance count; outer pie ring), and b) the types of substance combinations consumed (substance type; inner pie ring). First, we combined doses of ethanol and fentanyl, resulting in 3 substance types (water, alcohol, and fentanyl) and 7 substance combinations (Figure 2a). When examining all days together, both male and female mice drank single substances the majority of the time but also displayed polysubstance use (Figure 2b), and there was no significant difference in the distribution of substance counts (outer ring) between male and female mice (χ2=0.81, p>0.05). Conversely, there was a significant difference in the distribution of substance combinations (inner ring) between male and female mice (χ2=25.91, p<0.001). Next we examined each day of use separately and potential differences over successive days in the SIP chambers (Figure 2c). We found no differences in the distribution of substance counts (outer ring) between male and female mice on each of the 7 days (day 1: χ2=0.02, p>0.05; day 2: χ2=2.77, p>0.05; day 3: χ2=1.93, p>0.05; day 4: χ2=2.62, p>0.05; day 5: χ2=4.32, p>0.05; day 6: χ2=1.4, p>0.05; day 7: χ2=2.22, p>0.05). Conversely, there was a significant difference in the distribution of substance combinations (inner ring) between male and female mice on each of the 7 days (day 1: χ2=28.26, p<0.00001; day 2: χ2=14.95, p<0.05; day 3: χ2=22.78, p<0.001; day 4: χ2=29.32, p<0.00001; day 5: χ2=33.89, p<0.00001; day 6: χ2=35.75, p<0.00001; day 7: χ2=44.15, p<0.00001). Likewise, we found significant differences in both male and female mice across time in the distribution of substance counts (outer ring) (female day 1 vs 7: χ2=18.1, p<0.001; male day 1 vs 7: χ2=11.9, p<0.01) and in the distribution of substance combinations (inner ring) (female day 1 vs 7: χ2=32.7, p<0.00001; male day 1 vs 7: χ2=14.8, p>0.05).

**Figure 2:**
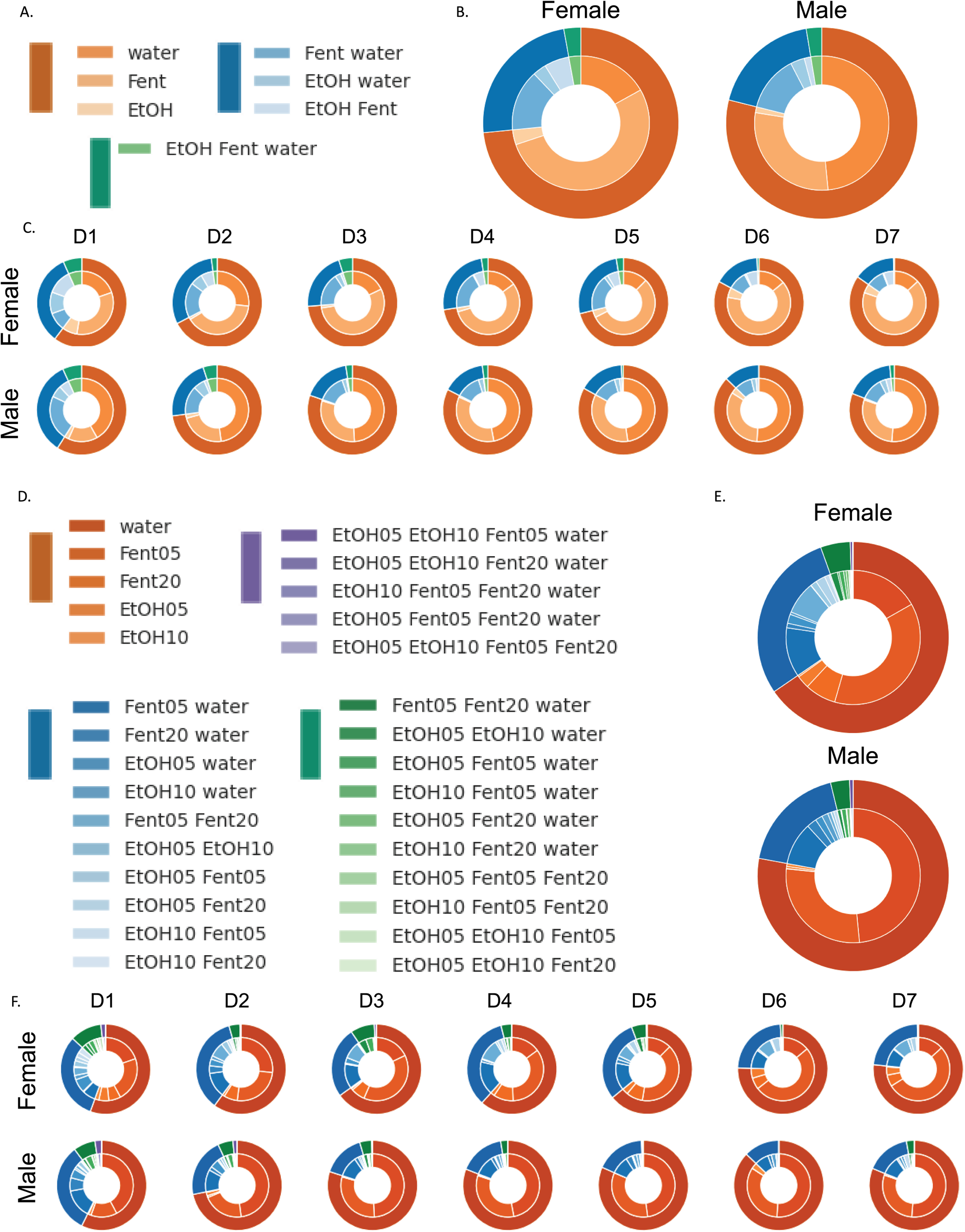
Polysubstance use in male and female mice. Polysubstance use was defined as intake of multiple substances within a 60 minute time frame. In each nested pie chart, the outer ring is the count of substance types consumed and the inner ring is the type of substance combination. A-C: Polysubstance use patterns with combined ethanol and fentanyl doses (3 substance types; 7 combinations). D-F: Polysubstance use patterns with individual ethanol and fentanyl doses (5 substance types; 30 combinations).

In a second set of visualizations, we examined doses of ethanol and fentanyl separately, resulting in 5 substance types and 30 substance combinations (Figure 2d-f). When examining all days together, both male and female mice drank single substances the majority of the time but also displayed polysubstance use (Figure 2e), and there was no significant difference in the distribution of substance counts (outer ring) between male and female mice (χ2=4.04, p>0.05). Likewise, there was no significant difference in the distribution of substance combinations (inner ring) between male and female mice (χ2=37.13, p>0.05). Next we examined each day of use separately and potential differences over successive days in the SIP chambers (Figure 2f). We found significant differences in the distribution of substance counts (outer ring) between male and female mice on days 4 and 5 (day 1: χ2=0.79, p>0.05; day 2: χ2=6.42, p>0.05; day 3: χ2=5.71, p>0.05; day 4: χ2=9.20, p<0.01; day 5: χ2=9.54, p<0.05; day 6: χ2=5.14, p>0.05; day 7: χ2=3.49, p>0.05). Likewise, there was a significant difference in the distribution of substance combinations (inner ring) between male and female mice on days 3-7 (day 1: χ2=41.79, p>0.05; day 2: χ2=23.65, p>0.05; day 3: χ2=40.39, p<0.01; day 4: χ2=45.45, p<0.001; day 5: χ2=48.95, p<0.001; day 6: χ2=57.52, p<0.00001; day 7: χ2=58.55, p<0.00001). Likewise, we found significant differences in both male and female mice across time in the distribution of substance counts (outer ring) (female day 1 vs 7: χ2=16.77, p<0.001; male day 1 vs 7: χ2=14.69, p<0.01) but only female mice significantly differed in the distribution of substance combinations from day 1 to 7 (inner ring) (female day 1 vs 7: χ2=42.24, p<0.05; male day 1 vs 7: χ2=27.43, p>0.05).

In addition to potential differences between male and female mice, we also hypothesized that baseline behavioral phenotypes might map on to subsequent polysubstance use profiles. One week prior to the start of housing in the SIP cages, mice were tested in the OFB and EZM. While male and female mice did not differ significantly in locomotor or anxiety-like metrics in the OFB (Figure 3a-f) or in the EZM (Figure 3g-l) (see Table 2 for statistical results), there was large amount of variability across animals, leading us to hypothesize that we could identify phenotypic sub-groups by using an unsupervised cluster-based approach. Given the high degree of collinearity across the 12 behavioral parameters collected from the OFB and EZM, we first performed a dimensionality reduction step using Principal Component Analysis (PCA) (Figure 3m-o). We then used the first three principal components (explaining over 75% of model variance) in a K-means cluster-based approach. Analysis of cluster stability supported a three-cluster solution (Table 3; Figure 3p-r). Using k=3, there is a non-significant trend for a different distribution of cluster assignment across male and female mice (Chi^2^ = 4.5, p=0.1) (Figure 3s). To determine whether OFB and EZM behavior differed across clusters, we assessed the 12 behavioral parameters when grouped by cluster assignment. Behavior across clusters differed significantly on all 12 parameters examined except for EZM open arm time (see Table 4 for statistical results) (Figure 3t-ae).

**Figure 3:**
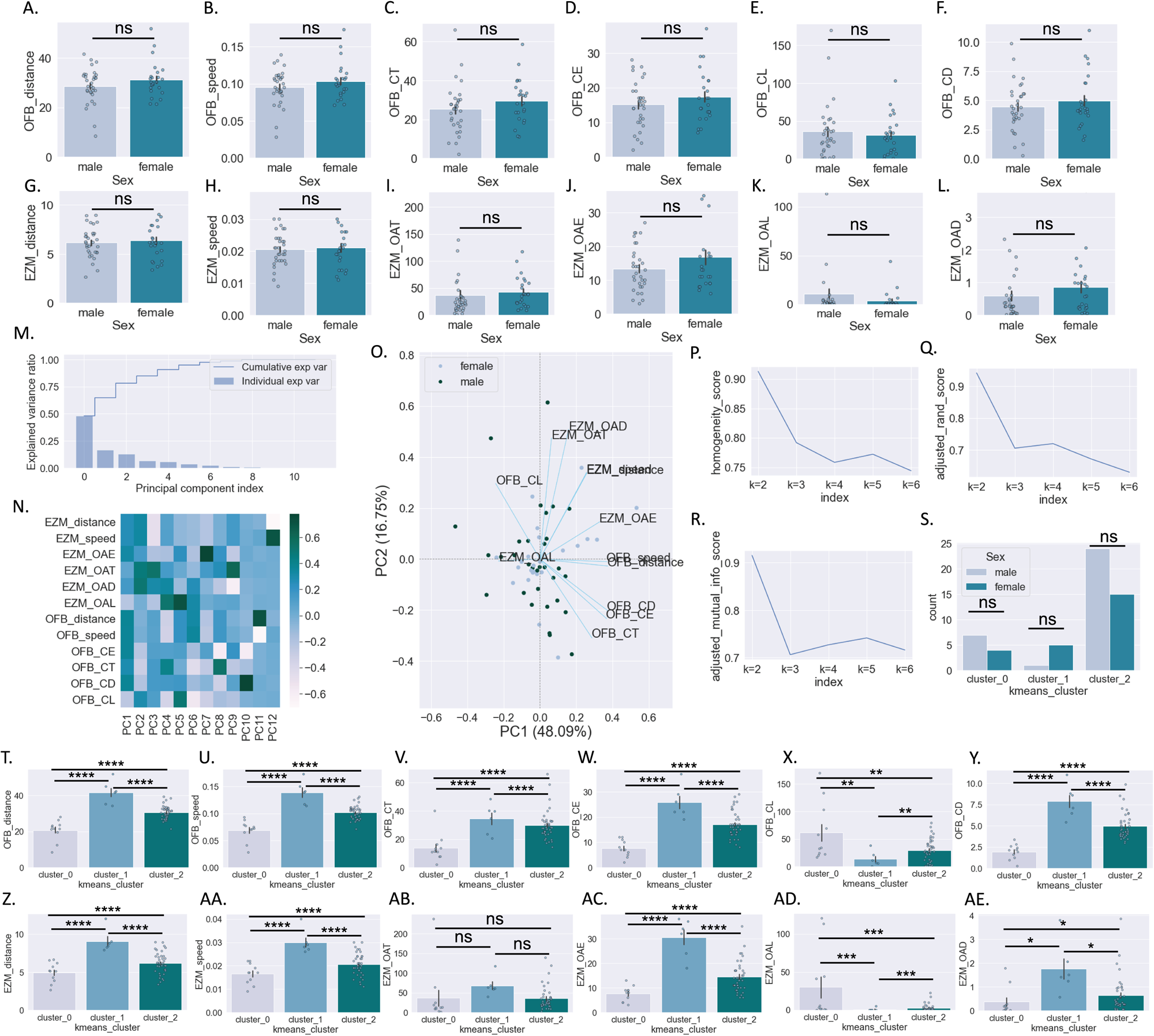
Baseline behavioral testing and cluster analysis. A-E : Behavioral parameters measured in the OFB in male and female mice. G-L: Behavioral parameters measured in the EZM in male and female mice. M-O. PCA dimensionality reduction of 12 behavioral parameters measured in OF and EZM: explained variance by PC (m). Heatmap of PC loadings by behavioral parameter (n). PCA biplot (o). P-Q: Unsupervised k-means clustering metrics using first three behavioral PCs. S: Behavioral cluster assignment by sex. T-Y: Behavioral parameters measured in the OFB by behavioral cluster. Z-AE: Behavioral parameters measured in the EZM behavioral cluster. Student’s t-test (a-l); Chi^2^ (s); One-way ANOVA *post hoc* BMCT (t-ae). **p ≤ 0.01, ****p ≤ 0.0001. Values represent mean ± SEM.

**Table 2:**
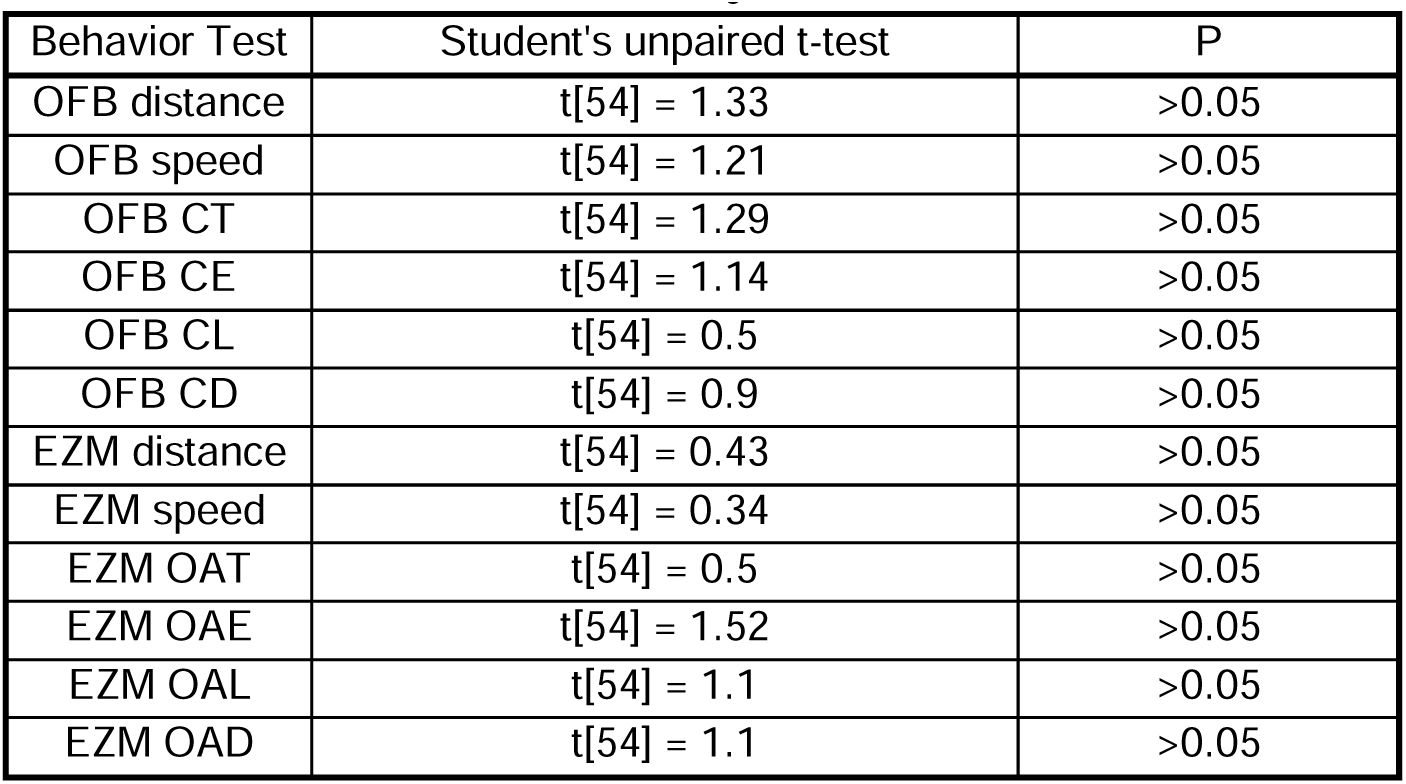
baseline behavioral data by sex.

| Behavior Test | Student's unpaired t-test | P |
| --- | --- | --- |
| OFB distance | t[54] = 1.33 | >0.05 |
| OFB speed | t[54] = 1.21 | >0.05 |
| OFB CT | t[54] = 1.29 | >0.05 |
| OFB CE | t[54] = 1.14 | >0.05 |
| OFB CL | t[54] = 0.5 | >0.05 |
| OFB CD | t[54] = 0.9 | >0.05 |
| EZM distance | t[54] = 0.43 | >0.05 |
| EZM speed | t[54] = 0.34 | >0.05 |
| EZM OAT | t[54] = 0.5 | >0.05 |
| EZM OAE | t[54] = 1.52 | >0.05 |
| EZM OAL | t[54] = 1.1 | >0.05 |
| EZM OAD | t[54] = 1.1 | >0.05 |

**Table 3:** cluster stability.

| k | homogeneity | completeness | V measure | adj. rand info. | adj. mutual info. |
| --- | --- | --- | --- | --- | --- |
| 3 | 0.79 | 0.67 | 0.72 | 0.71 | 0.71 |
| 4 | 0.76 | 0.74 | 0.75 | 0.72 | 0.73 |
| 5 | 0.77 | 0.77 | 0.77 | 0.67 | 0.74 |
| 6 | 0.74 | 0.77 | 0.76 | 0.63 | 0.72 |

**Table 4:** baseline behavioral data by cluster.

| Behavior Test | one-way ANOVA | P |
| --- | --- | --- |
| OFB distance | F[2,53] = 44.1 | **<.0001** |
| OFB speed | F[2,53] = 42.7 | **<.0001** |
| OFB CT | F[2,53] = 11.8 | **<.0001** |
| OFB CE | F[2,53] = 26.8 | **<.0001** |
| OFB CL | F[2,53] = 7.0 | **<.01** |
| OFB CD | F[2,53] = 32.9 | **<.0001** |
| EZM distance | F[2,53] = 16.6 | **<.0001** |
| EZM speed | F[2,53] = 15.7 | **<.0001** |
| EZM OAT | F[2,53] = 1.6 | >0.05 |
| EZM OAE | F[2,53] = 28.3 | **<.0001** |
| EZM OAL | F[2,53] = 8.2 | **<.001** |
| EZM OAD | F[2,53] = 6.7 | **<.01** |

Using these three behavioral phenotype clusters, data was first examined by grouping all substances together to assess general differences in the amount of time spent in the drinking chambers and total intake by cluster (Supp Figure 2). We found no significant differences across clusters in the time spent in the drinking chambers than male mice (Supp Figure 2a-e) nor in total amount of fluid consumed (Supp Figure 2f-j).

Next we examined potential differences in fluid intake across behavioral clusters for each substance separately (Figure 4). When summarized across all days in the SIP system, there was a significant interaction between Cluster and Substance and main effect of Substance type but no main effect of Cluster (two-way RM ANOVA: interaction effect F[8,212]=2.55, p<0.05, main effect Cluster F[2,53]=0.85, p>0.05, main effect Substance F[4,212]=23.5, p>0.0001; Benjamini/Hochberg FDR correction) (Figure 4a). Potential differences in amount consumed for each cluster across substances and light/dark cycle was examined using a separate two-way ANOVA for light vs. dark cycle (Dark cycle: two-way RM ANOVA: interaction effect F[8,212]=2.7, p<0.01, main effect Cluster F[2,53]=0.52, p>0.05, main effect Substance F[4,212]=24.4, p<0.0001; BMCT post hoc; Light cycle: two-way RM ANOVA: interaction effect F[8,2122]=1.67, p<0.05, main effect Cluster F[2,53]=1.13, p>0.05, main effect Substance F[4,212]=16.3, p<0.0001; BMCT post hoc). Next, we examined potential differences across days and behavioral cluster assignment for each substance separately (Figure 4c; see Table 5 for statistical results). Heat maps depicting the average amount of each substance consumed for each cluster across days and zeitgeber time are shown in Figure 4d.

**Figure 4:**
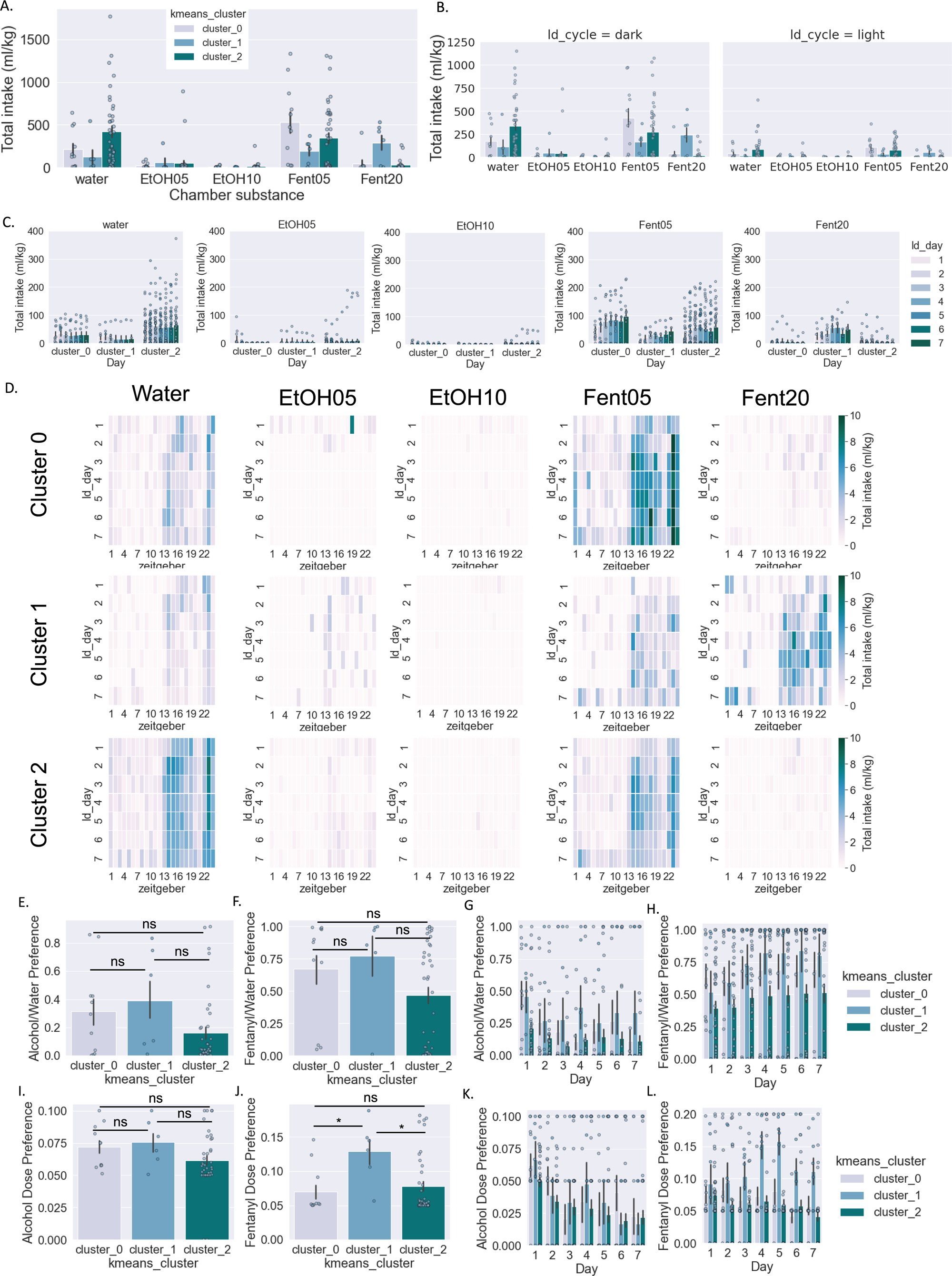
Consumption of individual substance/dose by cluster. A-C: Amount of liquid consumed by cluster (normalized to body weight) for each individual substance/dose combination in total (a), across light/dark cycle (b), and across days (c). D. Heatmap of amount consumed for each substance for individual mice. E. Heatmap of amount consumed for each substance by cluster across days and Zeitgeber time. F-I: Alcohol preference over water (f,g) and alcohol dose preference (h,i). J-M: Fentanyl preference over water (j,k) and fentanyl dose preference (l,m). Two-way RM ANOVA *post hoc* BMCT (a-c, g,i,k,m). Student’s t-test (f,h,j,l). *p ≤ 0.05, **p ≤ 0.01. Values represent mean ± SEM.

**Table 5:**
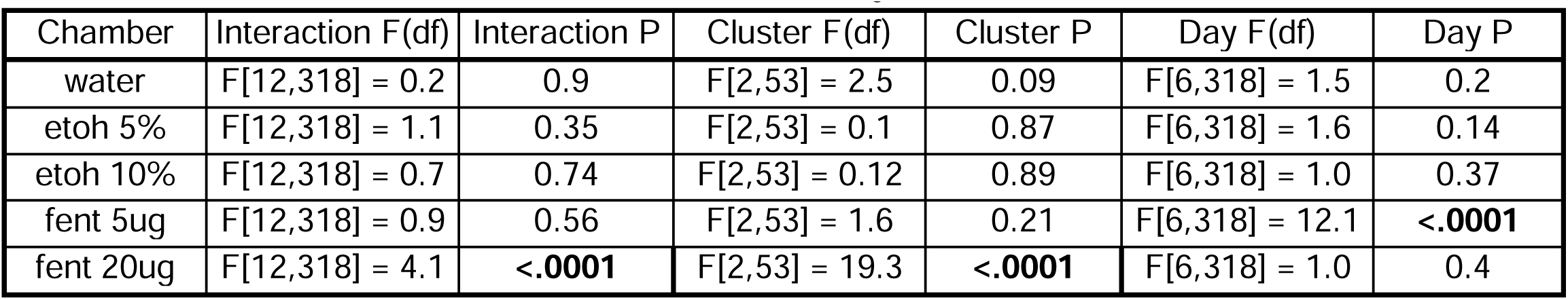
amount consumed across substance and day.

| Chamber | Interaction F(df) | Interaction P | Cluster F(df) | Cluster P | Day F(df) | Day P |
| --- | --- | --- | --- | --- | --- | --- |
| water | F[12,318] = 0.2 | 0.9 | F[2,53] = 2.5 | 0.09 | F[6,318] = 1.5 | 0.2 |
| etoh 5% | F[12,318] = 1.1 | 0.35 | F[2,53] = 0.1 | 0.87 | F[6,318] = 1.6 | 0.14 |
| etoh 10% | F[12,318] = 0.7 | 0.74 | F[2,53] = 0.12 | 0.89 | F[6,318] = 1.0 | 0.37 |
| fent 5ug | F[12,318] = 0.9 | 0.56 | F[2,53] = 1.6 | 0.21 | F[6,318] = 12.1 | **<.0001** |
| fent 20ug | F[12,318] = 4.1 | **<.0001** | F[2,53] = 19.3 | **<.0001** | F[6,318] = 1.0 | 0.4 |

When examining total intake, neither preference for alcohol nor fentanyl was significantly different across clusters (one-way ANOVA: F[2,53]=2.6, p>0.05, Figure 4e; one-way ANOVA: F[2,53]=2.2, p>0.05, Figure 4f). When examined across days, there was a significant main effect of Day but not Cluster or interaction effect for both ethanol and fentanyl preference (Figure 4g-h; see Table 6 for statistical results). Finally, we examined potential differences across behavioral clusters in the dose preference for alcohol and fentanyl (Figure 4i-l). When examining total intake, dose preference for alcohol was not significantly different across clusters (one-way ANOVA: F[2,53]=1.9, p>0.05) (Figure 4i). Conversely, dose preference for fentanyl was significantly different across clusters (one-way ANOVA: F[2,53]=4.4, p<0.05) (Figure 4j). When examined across days, there was only a significant main effect of Day for ethanol dose preference, while there were significant main effects of Day and Cluster for fentanyl dose preference (Figure 4k-l; see Table 7 for statistical results).

**Table 6:** ethanol/water and fentanyl/water preference across days.

| Substance | Interaction F(df) | Interaction P | Cluster F(df) | Cluster P | Day F(df) | Day P |
| --- | --- | --- | --- | --- | --- | --- |
| Alcohol | F[12,318] = 0.8 | >0.05 | F[2,53] = 2.3 | >0.05 | F[6,318] = 3.2 | **<.01** |
| Fentanyl | F[12,318] = 0.7 | >0.05 | F[2,53] = 1.9 | >0.05 | F[6,318] = 5.7 | **<.001** |

**Table 7:** ethanol and fentanyl dose preference across days.

| Substance | Interaction F(df) | Interaction P | Cluster F(df) | Cluster P | Day F(df) | Day P |
| --- | --- | --- | --- | --- | --- | --- |
| Alcohol | F[12,318] = 0.6 | >0.05 | F[2,53] = 0.7 | >0.05 | F[6,318] = 6.2 | **<.001** |
| Fentanyl | F[12,318] = 1.7 | >0.05 | F[2,53] = 5.6 | **<.01** | F[6,318] = 2.7 | **<0.05** |

In addition to examining each substance separately, we also investigated potential polysubstance use across behavioral clusters (Figure 5). First, we combined doses of ethanol and fentanyl, resulting in 3 substance types (water, alcohol, and fentanyl) and 7 substance combinations (Figure 5a). When examining all days together, all three clusters drank single substances the majority of the time but also displayed polysubstance use (Figure 5b), and there was no significant difference in the distribution of substance counts (outer ring) between clusters (χ2=1.25, p>0.05). Conversely, there was a significant difference in the distribution of substance combinations (inner ring) across the three clusters (χ2=25.38, p<0.05). Next we examined each day of use separately and potential differences over successive days in the SIP chambers (Figure 5c). We found significant differences in the distribution of substance counts (outer ring) between clusters only on day 3 (day 1: χ2=4.2, p>0.05; day 2: χ2=8.22, p>0.05; day 3: χ2=10.28, p>0.05; day 4: χ2=1.95, p>0.05; day 5: χ2=3.11, p>0.05; day 6: χ2=0.17, p>0.05; day 7: χ2=5.67, p>0.05). Conversely, there was a significant difference in the distribution of substance combinations (inner ring) between clusters on all 7 days (day 1: χ2=32.6, p>0.01; day 2: χ2=38.67, p<0.001; day 3: χ2=25.65, p<0.05; day 4: χ2=45.5, p>0.000001; day 5: χ2=33.41, p<0.001; day 6: χ2=42.7, p>0.000001; day 7: χ2=52.37, p<0.000001). Likewise, we found significant differences across clusters and time in the distribution of substance counts (outer ring) (cluster 1 day 1 vs 7: χ2=22.94, p<0.00001; cluster 2 day 1 vs 7: χ2=16.84, p<0.001; cluster 3 day 1 vs 7: χ2=11.48, p<0.01) and in the distribution of substance combinations (inner ring) (cluster 1 day 1 vs 7: χ2=35.23 p<0.000001; cluster 2 day 1 vs 7: χ2=57.0, p<0.000001; cluster 3 day 1 vs 7: χ2=14.78, p<0.05).

**Figure 5:**
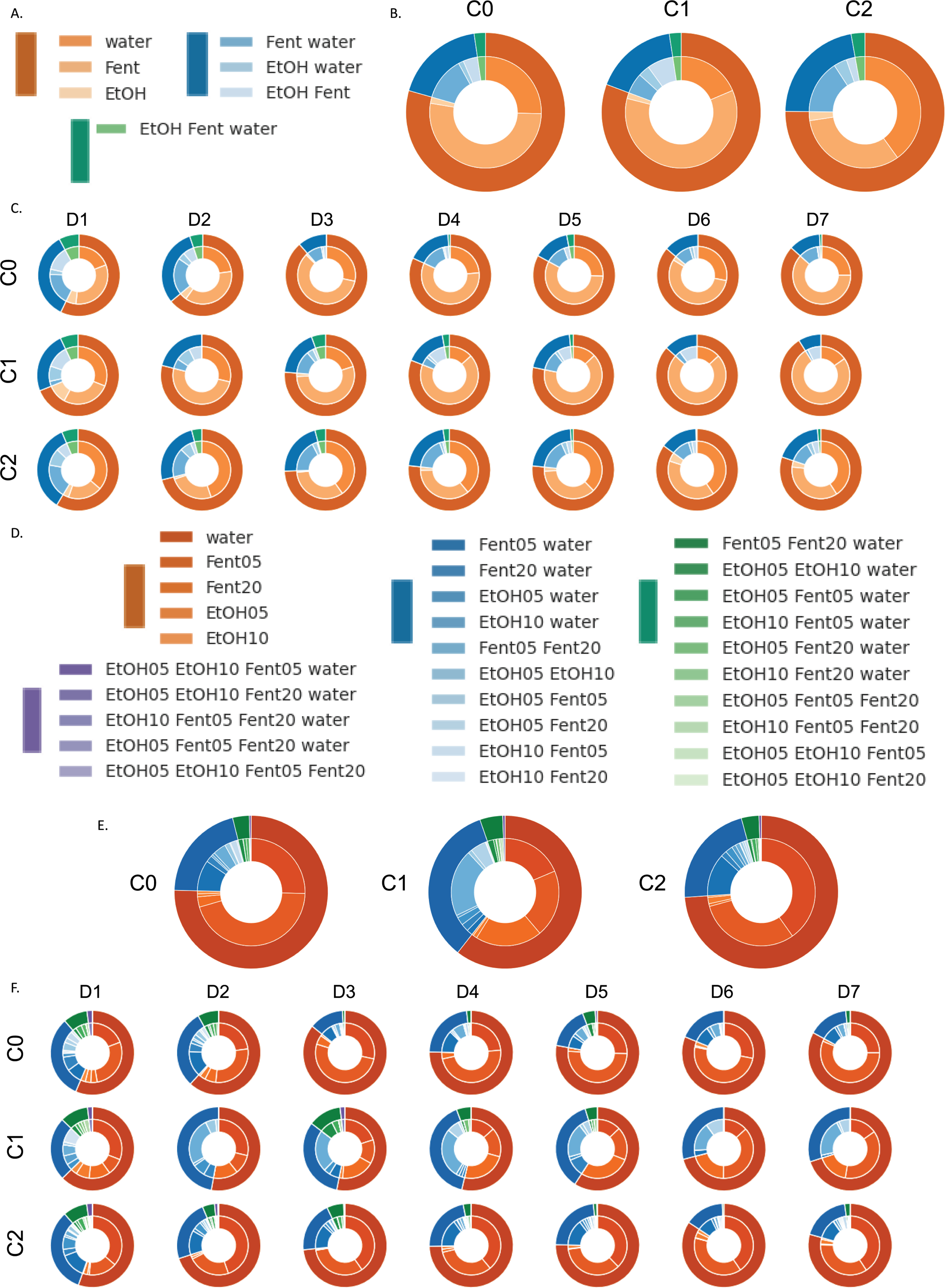
Polysubstance use by behavioral cluster. Polysubstance use was defined as intake of multiple substances within a 60 minute time frame. In each nested pie chart, the outer ring is the count of substance types consumed and the inner ring is the type of substance combination. A-C: Polysubstance use patterns with combined ethanol and fentanyl doses (3 substance types; 7 combinations). D-F: Polysubstance use patterns with individual ethanol and fentanyl doses (5 substance types; 30 combinations).

In a second set of visualizations, we examined doses of ethanol and fentanyl separately, resulting in 5 substance types and 30 substance combinations (Figure 5d-f). When examining all days together, all three clusters drank single substances the majority of the time but also displayed polysubstance use (Figure 5e), and there was no significant difference in the distribution of substance counts (outer ring) across clusters (χ2=6.49, p>0.05). Conversely, there was a significant difference in the distribution of substance combinations (inner ring) across clusters (χ2=93.97, p<0.001). Next we examined each day of use separately and potential differences over successive days in the SIP chambers (Figure 5f). We found significant differences in the distribution of substance counts (outer ring) across clusters on days 2-5 (day 1: χ2=2.17, p>0.05; day 2: χ2=22.22, p<0.01; day 3: χ2=28.71, p<0.000001; day 4: χ2=14.38 p<0.01; day 5: χ2=14.24, p<0.05; day 6: χ2=7.44, p>0.05; day 7: χ2=8.53, p>0.05). Conversely, there was a significant difference in the distribution of substance combinations (inner ring) across clusters on all 7 days of intake (day 1: χ2=111.92, p<0.000001; day 2: χ2=114.38, p<0.000001; day 3: χ2=120.72, p<0.000001; day 4: χ2=134.85, p<0.000001; day 5: χ2=146.48, p<0.000001; day 6: χ2=107.89, p<0.000001; day 7: χ2=112.48, p<0.000001). Likewise, we found significant differences across time in the distribution of substance counts for each cluster (outer ring) (cluster 0 day 1 vs 7: χ2=18.14, p<0.001; cluster 1 day 1 vs 7: χ2=13.52, p<0.01; cluster 2 day 1 vs 7: χ2=15.2, p<0.01) and in the distribution of substance combinations (inner ring) for clusters 0 and 1 but not cluster 2 (cluster 0 day 1 vs 7: χ2=42.7, p<0.001; cluster 1 day 1 vs 7: χ2=77.7, p<0.000001; cluster 2 day 1 vs 7: χ2=26.32, p<0.05).

## Discussion

This study aimed to better understand how individual differences influence alcohol and opioid intake and polysubstance use in male and female mice. We identified multiple parameters related to drinking activity that differed according to sex. We also uncovered three discrete clusters of mice based on behavioral phenotypes that had unique drinking patterns. Together our results demonstrate the utility of studying polysubstance use in group housed mice and support the overarching notion that baseline behavioral phenotypes relate to substance use and preference patterns.

The first outcome that we measured was activity level, determined by number of visits to and time spent in the drinking chambers (registered by RFID sensor). While the number of visits and time spent in the drinking chambers is an imperfect measure of activity, it gives an initial baseline to build from. Both male and female mice decreased time spent in the chambers across the seven days in the SIP system, but female mice spent more time in the drinking chambers each day. This agrees with previous rodent studies that found increased locomotion in female rodents compared to males after chronic alcohol, fentanyl, or morphine administration^24–27^. It is unclear why differences in locomotion exist between male and female rodents following alcohol and/or opioid consumption. One possible explanation could be differences in metabolism and how these substances physiologically affect males and females, or potentially differences in the rewarding or aversive neural properties of a substance. Importantly, we did not track estrous cycle in the female mice. While changes in estrous cycle could potentially influence the reinforcing effects of fentanyl, previous studies have shown that estrous cycle likely does not impact locomotor behavior^28,29^.

When looking at intake across the five available substances (water, 5% and 10% alcohol, 5ug and 20ug fentanyl), there were sex differences in substance intake pattern and preference. On average, male mice consumed the most water, followed closely by 5ug fentanyl, small amounts of 5% and 10% ethanol, and the lowest volume of 20ug fentanyl. The highest total intake for female mice was 5ug fentanyl, then water, closely followed by 20ug fentanyl, 5% ethanol, and the smallest volume of 10% ethanol. Males had a slight preference for alcohol over water and a moderate preference for fentanyl over water, while females had a moderate preference for alcohol and a strong preference for fentanyl over water. In females, the preference for alcohol over water decreased over time, but fentanyl preference escalated over time. Fentanyl preference remained generally consistent for the male mice. There were no statistical differences in dose preference between male and female mice.

Our results generally corroborate trends seen previously. Female mice tend to consume higher amounts of ethanol^24,30^ and fentanyl^31–33^ relative to their body weight compared to male mice. Another study found that female rats drank larger volumes of a 5% dose of ethanol compared to male mice, as well as compared to other higher doses of ethanol, showing the importance of including multiple doses of substances^24^. There is also evidence in both human and rodent studies that females will escalate from initial and moderate substance consumption to disordered use or addiction more quickly than males^32,34^, which mirrors what we saw with the female mice escalating fentanyl preference during the seven days.

We also found significant differences in polysubstance patterns (here defined as the intake of 2 or more substances within the same hour) in relation to both sex and across days in the SIP chambers. In general, single substance use predominated and polysubstance use was highest on day 1 and decreased over successive days of intake. Female mice displayed increased and more sustained polysubstance use than male mice. Currently, there are published studies reporting opioid followed by alcohol use in rats (sequential polysubstance use) (Wilkinson et al., 2023; Wilkinson et al., 2025), and one study investigated concurrent oxycodone and alcohol self-administration but did not report on polysubstance use patterns (Amico et al., 2022). The narrowing of polysubstance use across successive days in the SIP chambers may represent an initial exploration phase where animals sample from multiple substances to a more ‘substance of choice’ phase.

One striking result from this study is the high variability in consumption, not only between mice but also across days within individual mice. The constant access and voluntary consumption model of the SIP system provides an abundance of data regarding the timing and dose preference patterns for each individual mouse. When looking to the clinical literature to uncover motivators underlying choice in substance use, it appears choice is often driven by stress-related experience, social environment, or personality traits such as impulsivity and maladaptive coping strategies^9,15,35–37^. To test this concept using our SIP system, we decided to assess locomotion and anxiety-like behaviors one week prior to housing in the SIP cages to investigate any correlations between behavior and substance use patterns.

Our initial examination revealed no significant sex differences on any of the twelve parameters in the open field (OFB) and elevated zero maze (EZM) tests. While behavioral tests have some degree of variability, typically female rodents show lower anxiety-like behaviors, with no sex differences in novelty-seeking behavior (although this can depend on estrous phase)^38,39^. Because there was a considerable amount of variability across mice in our study, we hypothesized that the range of behavioral profiles might map on polysubstance use patterns. After dimensionality reduction and an unsupervised clustering analysis based on the 12 behavioral parameters, three distinct groups of mice were revealed. The composition of male and female mice in cluster zero had 7 males and 4 females, cluster one had 5 females and 1 male, and cluster two had 24 males and 15 females; this distribution was trending but non-significant when tested statistically. The clusters did statistically differ in 11 of 12 behavioral parameters (all except EZM open arm time) which suggests we identified three distinct behaviorally phenotypic subgroups. Cluster 0 was defined by higher anxiety-like behaviors, including less distance traveled in the center of the OFB and in the open arms of the EZM and longest latency to enter the center area/open arms. Cluster 1 had the longest time spent in the center of the OFB and open arms of the EZM, and shortest latency to enter the center area/open arms, suggesting lower anxiety-like behavior.

Finally, we projected the three clusters onto the substance consumption data. Although this would not prove a causal relationship between behavioral phenotypes and polysubstance use patterns, it certainly provides beneficial insight and highlights predictive ability. There were meaningful differences in consumption patterns between the three clusters, with cluster 0 drinking a high amount of 5% fentanyl and a moderate amount of water; cluster 1 consuming a high amount of 20% fentanyl and a moderate amount of 5% fentanyl; and cluster 2 consuming a high amount of water, moderate 5% fentanyl and small amount of 5% ethanol. There were no significant differences between clusters for ethanol or fentanyl preference over water, or for alcohol dose preference, but fentanyl dose preference was higher for cluster 1 compared to clusters 0 and 2 and increased over the course of the seven days of substance access. Polysubstance use patterns also differed across behavioral clusters, with cluster 1 exhibiting the highest and most consistent polysubstance use patterns.

Taken all together, it appears that cluster 1 consists of majority female mice, shows lower anxiety-like behavior, and preferentially consumes a higher dose of fentanyl. Previous studies have found mixed results relating anxiety-like and novelty-seeking behaviors with higher opioid consumption^21^. In our study, Cluster 1 showed more exploratory and less anxious behavior and the highest consumption of fentanyl. Surprisingly, the cluster with the highest anxiety-like behavior (cluster 0) did not have the highest preference for ethanol, as has been shown before in the literature^17–19^. This could be because the mice had access to fentanyl in addition to the ethanol, the 24-hour access of the alcohol, the concentration of ethanol, or because there were no stressors prior to substance availability.

To our knowledge, there are only three other studies that consist of simultaneous or sequential (respectively) voluntary administration of an opioid and alcohol^5,40,41^. One study (Amico et al., 2022) examined alcohol and oxycodone co-use but did not report on specific polysubstance use patterns (but did find that female rats consumed more alcohol and oxycodone than male rats). The other two studies investigated oxycodone self-administration directly followed by homecage access to alcohol. In line with our research, Wilkinson et al., also found that male and female rats with access to oxycodone consumed less alcohol than rats that only had access to sucrose. Neither study conducted behavioral testing before alcohol or opioid administration. Likewise, Wilkinson et al., 2025 reported that elevated baseline anxiety-like behavior was accompanied by redacted alcohol intake, in line with our current results. While there are some meaningful differences that prevent direct comparison between these studies and our experiments here, a main takeaway is the persistent existence of sex differences in polysubstance use and behavioral profiles across a variety of housing conditions and access paradigms.

Our results provide an initial characterization of some of the fundamental parameters surrounding polysubstance use in a preclinical model, and the interpretation is constrained by the scope of the experiment. We began with seven days in the SIP cages, and while we observed escalation of consumption, changes in dose preference, and a narrowing of polysubstance use patterns, a longer experimental timeline will be critical to understand the transition from casual substance use to development of an SUD-like phenotype. We provided continuous access to both alcohol and fentanyl, and an intermittent access paradigm may reveal different patterns of use. We relied on drinking chamber visits to determine activity, which could not accurately reflect total locomotion. The addition of a stressor, or period of extinction/deprivation of a substance would also help improve our understanding of drug seeking and motivations for consumption. Age of first exposure is known to have significant implications for future substance consumption and behavioral and biological outcomes; so inclusion of animal models across the lifespan is important as well^4,24,42,43^. Future studies should investigate the mechanisms underlying drug metabolism and pharmacology and how it affects other related behaviors, including sex differences. Physiological measures and biomarkers could play an important role in predicting future substance consumption patterns, consequences of substance use, and treatment outcomes.

The SIP system provides an enriched social environment and voluntary consumption of multiple substances, and the possibilities for future studies using the SIP system are nearly unlimited. It offers the opportunity to continue interrogating the role of sex differences in substance use. It is pertinent to acknowledge that our preclinical models do have limitations in uncovering the multi-faceted and societal-driven motivations to consume substances that are cited in clinical studies, but some indicators such as anxiety-like behaviors and stress responses are preserved across species. These basic behaviors may help us to reveal critical factors that influence substance use. Overall, we hope this study underscores the need for more preclinical research on polysubstance use to better understand the patterns of consumption, treatment outcomes, and novel therapeutic strategies.

## Supporting information

supp results

## ABBREVIATIONS

AUD: Alcohol Use Disorder
BMCT: Bonferroni Multiple Comparison Test
CD: Center Distance
CE: Center Entries
CL: Center Latency
CT: Center Time
EtOH: Ethanol
EZM: Elevated Zero Maze
Fent: Fentanyl
OAD: Open Arm Distance
OAE: Open Arm Entries
OAL: Open Arm Latency
OAT: Open Arm Time
OFB: Open Field Box
OUD: Opioid Use Disorder
PC: Principal Component
SIP: Socially Integrated Polysubstance
SUD: Substance Use Disorder

## DISCLAIMER

The views expressed in this scientific presentation are those of the author(s) and do not reflect the official policy or position of the U.S. government or Department of Veteran Affairs.

## DECLARATIONS

### Ethics approval and consent to participate

All animal experiments were conducted in accordance with Association for Assessment and Accreditation of Laboratory Animal Care guidelines and were approved by the VA Puget Sound Institutional Animal Care and Use Committee.

### Consent for publication

Not applicable.

### Availability of data and materials

The data in this study are available from the corresponding author upon reasonable request.

### Competing interests

The authors declare that the research was conducted in the absence of any commercial or financial relationships that could be construed as a potential conflict of interest.

## Funding

This work was supported by grants from NIAAA Training Grant 5T32AA007455 (MP), NIDA Training Grant 2T32DA007278-26 (BMB), UW NAPE Summer Undergraduate Research Program NIDA DA048736 (KW), UW NAPE Pilot Program NIDA DA048736 (AGS), and Department of Veteran Affairs (VA) Basic Laboratory Research and Development (BLR&D) Merit Award BX005582 (AGS).

## Authors’ contributions

The work presented here was carried out in collaboration among all authors. MP, KW, BB, and AGS contributed to conception and design of the study. MP, ZCW, ABK, KW, SJL, ES, RN, BB, and AGS collected and analyzed data. MP and AGS wrote the first draft of the manuscript. All authors contributed to manuscript revision, read, and approved the final manuscript.

## Acknowledgements

We would like to thank Scott Ng Evans, Traci J. Weber, Cindy Pekow, DVM, Kari Koszdin, DVM, and Lena Strait-Bodey for technical assistance and veterinary care.

---

**Supp Figure 1: Activity and consumption in combined drinking chambers by sex**

A-E : Time spent in the drinking chambers in male and female mice in total (a), across days (b), across light/dark cycle (c), and across Zeitgeber time (d); heatmaps shown in e. F-J: Amount of liquid consumed by male and female mice (normalized to body weight) in total (f), across days (g), across light/dark cycle (h), and across Zeitgeber time (i); heatmaps shown in j. Student’s t-test (a,f); Two-way RM ANOVA *post hoc* BMCT (b-d. g-h). **p ≤ 0.01, ****p ≤ 0.0001. Values represent mean ± SEM.

**Supp Figure 2: Activity and consumption in combined drinking chambers by cluster**

A-E: Time spent in the drinking chambers for each cluster in total (a), across days (b), across light/dark cycle (c), and across Zeitgeber time (d); heatmaps shown in e. F-J: Amount of liquid consumed by mice in each cluster (normalized to body weight) in total (f), across days (g), across light/dark cycle (h), and across Zeitgeber time (i); heatmaps shown in j. Student’s t-test (a,f); Two-way RM ANOVA *post hoc* BMCT (b-d. g-h). **p ≤ 0.01, ****p ≤ 0.0001. Values represent mean ± SEM.

## Notes

### Competing Interest Statement

The authors have declared no competing interest.

### Summary of Updates

Addition of polysubstance intake pattern analysis.

