## Supplementary material for "Anxiety and risk-taking behavior maps onto opioid and alcohol polysubstance consumption patterns in male and female mice": supp results

Supp Figure 1:

Visit summary data by sex: Female mice spent more time than male mice in the drinking chambers in total (Student’s unpaired t-test, t[54]=5.12, p<0.0001) (Supp. Figure 1a) and when analyzed across days (two-way RM ANOVA: interaction effect F[6,324]=1.47, p>0.05, main effect Sex F[1,54]=26.3, p<0.0001, main effect Day F[6,324]=5.7, p>0.0001; Bonferroni Multiple Comparison Test (BMCT) post hoc) (Supp. Figure 1b), and both sexes decreased time spent in the chambers as days in the SIP system progressed. Female mice spent more time in the drinking chambers specifically during the dark cycle (with both sexes showing decreased time spent during the light vs. dark cycle) (two-way RM ANOVA: interaction effect F[1,54]=24.9, p<0.001, main effect Sex F[1,54]=26.3, p<0.0001, main effect Cycle F[1,54]=206.3, p>0.0001; BMCT post hoc) (Supp. Figure 1c). Likewise, female mice spent more time in the drinking chambers during all hours of the dark cycle and some hours of the light cycle (two-way RM ANOVA: interaction effect F[23,1242]=10.9, p<0.001, main effect Sex F[1,54]=26.3, p<0.0001, main effect Zeitgeber F[23,1242]=121.2, p>0.0001; BMCT post hoc) (Supp. Figure 1d). Heat maps depicting average time spent in the drinking chambers for male and female mice across days and zeitgeber time are shown in Supp. Figure 1e.

Drinking summary data by sex: Female and male mice did not differ in the total amount of liquid consumed (Student’s unpaired t-test, t[54]=1.1, p>0.05) (Supp. Figure 1f). When analyzed across days, there was a significant interaction effect but no main effect of sex (two-way RM ANOVA: interaction effect F[6,324]=6.4, p<0.001, main effect Sex F[1,54]=1.2, p>0.05, main effect Day F[6,324]=3.8, p>0.0001; BMCT post hoc) (Supp. Figure 1g), with both sexes increasing amount consumed as days in the SIP system progressed. When examined by light/dark cycle, there was a significant interaction effect but no main effect of sex (with both sexes showing a decrease in amount consumed during the light vs. dark cycle) (two-way RM ANOVA: interaction effect F[1,54]=5.6, p<0.02, main effect Sex F[1,54]=1.2, p>0.05, main effect Cycle F[1,54]=356.7, p>0.0001; BMCT post hoc) (Supp. Figure 1h). Likewise, when examined by zeitgeber time, there was a significant interaction effect with female mice consuming more liquid during all hours of the dark cycle and some hours of the light cycle (two-way RM ANOVA: interaction effect F[23,1242]=3.1, p<0.001, main effect Sex F[1,54]=1.2, p>0.05, main effect Zeitgeber F[23,1242]=3.0, p>0.0001; BMCT post hoc) (Supp. Figure 1i). Heat maps depicting the average amount consumed in drinking chambers for male and female mice across days and zeitgeber time are shown in Supp. Figure 1j.

Supp Figure 2:

Visit summary data by behavioral cluster: There was no significant difference across clusters for total time spent in the drinking chambers (one-way ANOVA: F[2,53]=1.22, p>0.05) (Supp. Figure 2a). When analyzed across days, there was a significant effect of Day but not Cluster (two-way RM ANOVA: interaction effect F[12,318]=1.62, p>0.05, main effect Cluster F[2,53]=1.2, p>0.05, main effect Day F[6,318]=5.7, p<0.0001; BMCT post hoc) (Supp. Figure 2b). Likewise, when examined by light/dark cycle, there was a significant effect of Cycle but not Cluster (with all clusters showing decreased time spent during the light vs. dark cycle) (two-way RM ANOVA: interaction effect F[2,53]=1.3, p>0.05, main effect Cluster F[2,53]=1.2, p>0.05, main effect Cycle F[1,53]=151.3, p<0.0001; BMCT post hoc) (Supp. Figure 2c). Finally, when examined by zeitgeber time, there was a significant effect of Time but not Cluster (two-way RM ANOVA: interaction effect F[46,1219]=1.2, p>0.05, main effect Cluster F[2,53]=1.2, p>0.05, main effect Time F[23,1219]=103.6, p<0.0001; BMCT post hoc) (Supp. Figure 2d). Heat maps depicting the average time spent in the drinking chambers for males and females across days and zeitgeber time are shown in Supp. Figure 2e.

Drinking summary data by behavioral cluster: There was no significant difference across clusters for total liquid consumed (one-way ANOVA: F[2,53]=0.84, p>0.05) (Supp. Figure 2f). When analyzed across days, there was a significant effect of Day but not Cluster (two-way RM ANOVA: interaction effect F[12,318]=0.43, p>0.05, main effect Cluster F[2,53]=0.84, p>0.05, main effect Day F[6,318]=8.7, p<0.0001; BMCT post hoc) (Supp. Figure 2g). Likewise, when examined by light/dark cycle, there was a significant effect of Cycle but not Cluster (with all clusters showing decreased time spent during the light vs. dark cycle) (two-way RM ANOVA: interaction effect F[2,53]=0.01, p>0.05, main effect Cluster F[2,53]=0.8, p>0.05, main effect Cycle F[1,53]=318.4, p<0.0001; BMCT post hoc) (Supp. Figure 2h). When examined by zeitgeber time, there was a significant effect of Time but not Cluster (two-way RM ANOVA: interaction effect F[46,1219]=0.8, p>0.05, main effect Cluster F[2,53]=0.9, p>0.05, main effect Time F[23,1219]=97.2, p<0.0001; BMCT post hoc) (Supp. Figure 2i). Heat maps depicting average time spent in the drinking chambers for male and female mice across days and zeitgeber time are shown in Supp. Figure 2j.
