## Supplementary figures and images for "Anxiety and risk-taking behavior maps onto opioid and alcohol polysubstance consumption patterns in male and female mice"

### sf1

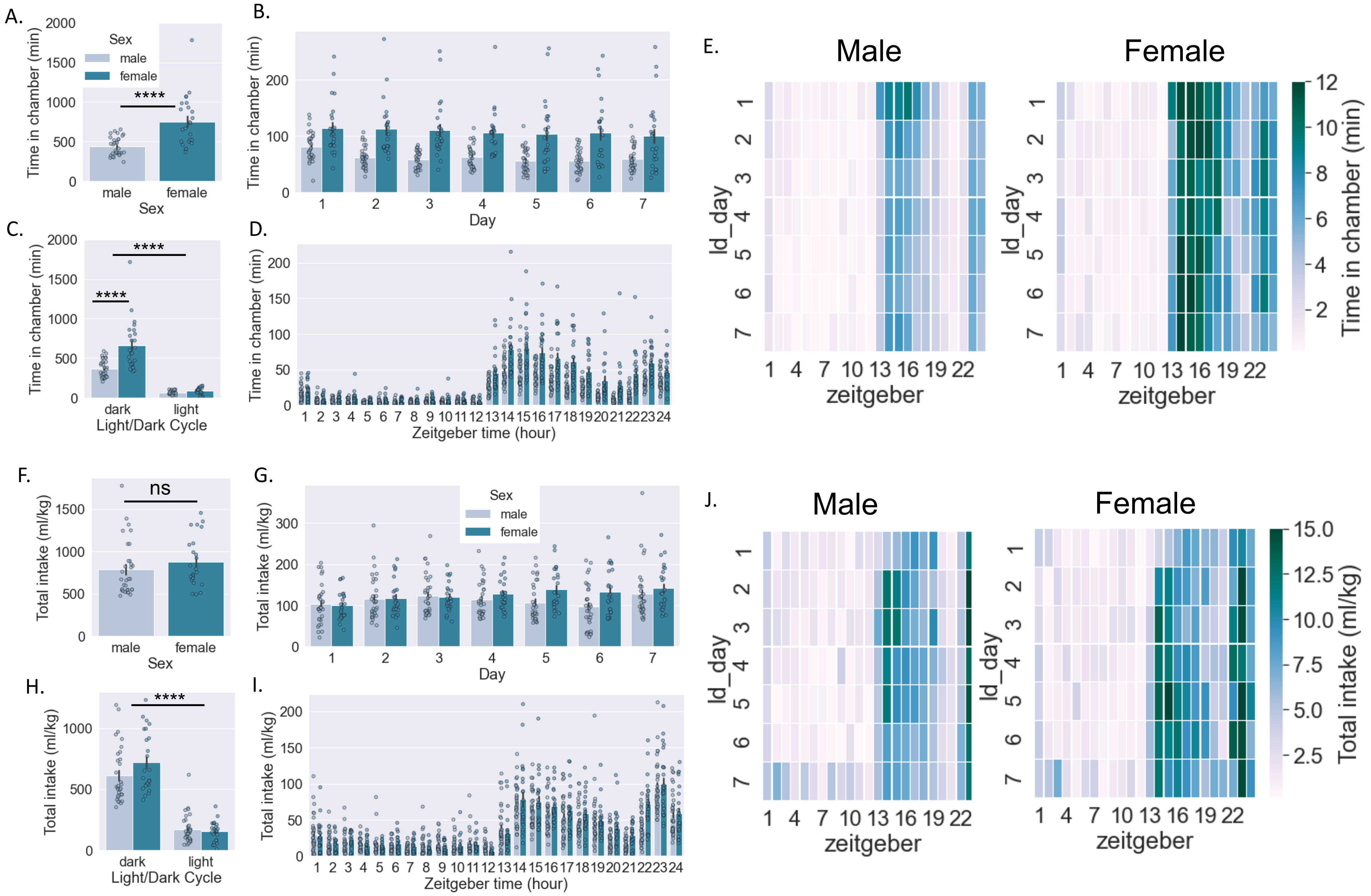

### sf2

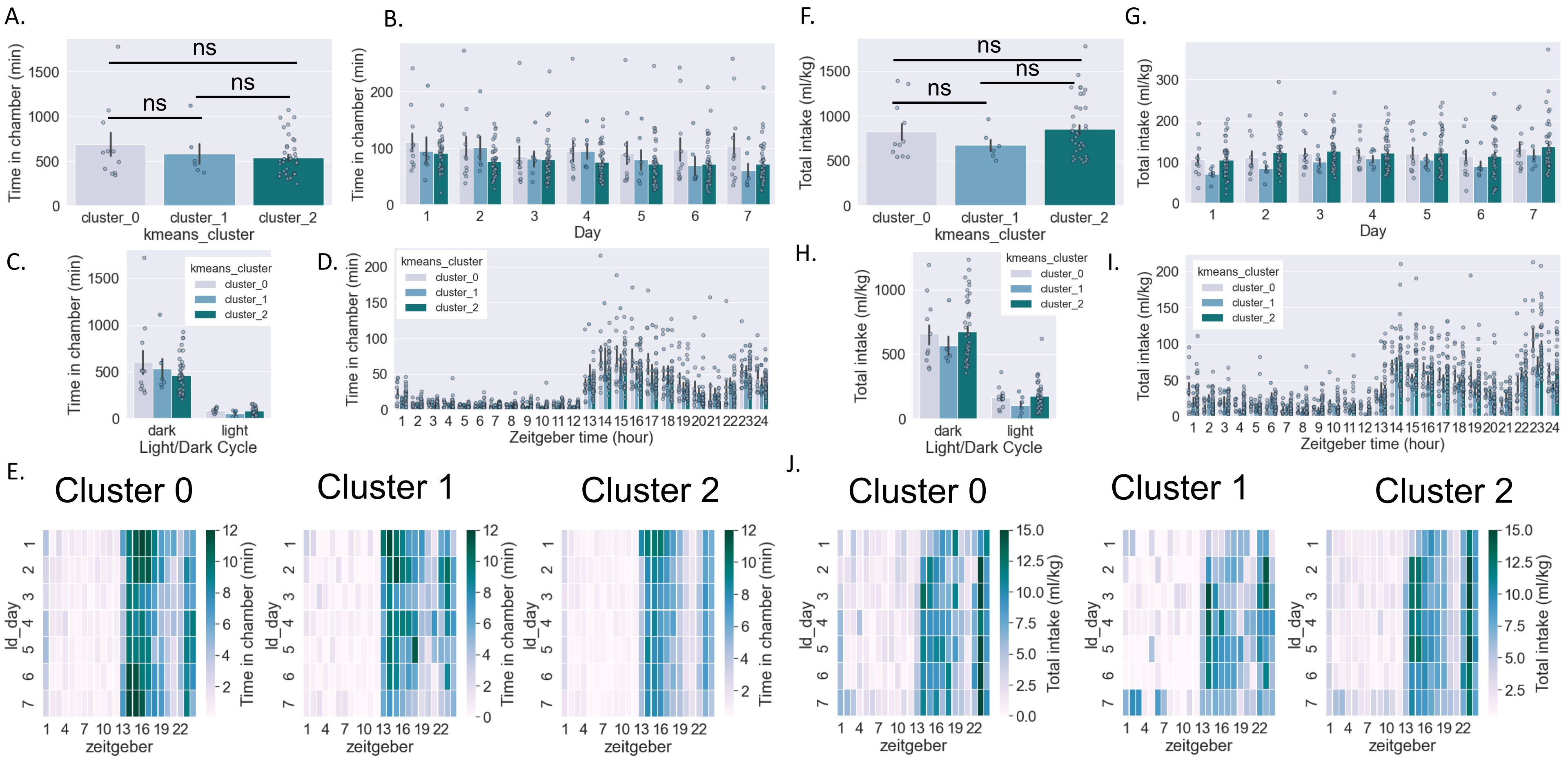
